# Sequencing of *α*-synuclein Intrinsically Disordered Protein in MoS_2_ Nanopores

**DOI:** 10.1101/2025.07.23.666249

**Authors:** Adrien Nicolaï, Patrice Delarue, Andreina Urquiola Hernandez, Patrick Senet

## Abstract

*α*-synuclein is an intrinsically disordered protein composed of 140 amino acids that adopts multiple conformations and is prone to aggregate into *β*-sheet-rich structures, which are hallmarks of Parkinson’s disease. Moreover, missense mutations in *α*-synuclein are related to familial forms of this neurodegenerative disorder, making their detection at the single-molecule level essential. Recently, protein sequencing using solid-state nanopores has emerged as a powerful, label-free approach for single-molecule sensing with high sensitivity. Atomically thin two-dimensional materials, such as MoS_2_, provide ideal platforms for sequencing applications due to their ultimate thinness and enhanced spatial resolution. However, protein sequencing using 2D materials remains challenging because of rapid translocation speeds, which limit the observation time per molecule. Here, we present extensive all-atom classical molecular dynamics simulations in explicit solvent of the full translocation of the wild-type *α*-synuclein protein and two pathogenic mutants, *i*.*e*. A30P and E46K, through single-layer MoS_2_ nanopores, for a total duration of more than 22 *µ*s. To the best of our knowledge, this work represents the first atomistic simulation of a full-length protein sequencing through a 2D solid-state nanopore. From the ionic current traces recorded during simulations, we characterized distinct blockade levels and bumps, as well as their dwell time along the protein sequence at multiple time and sequence length scales. For instance, we provided the sequence motifs that show some particular patterns in the data. Furthermore, we analyzed the volume properties of amino acids inside the pore and identified characteristic blockade fingerprints differentiating the wild-type from the mutant proteins. These pioneering results pave the way for future experimental studies, offering a roadmap for validating 2D nanopore-based protein sequencing and biomarker detection with single-molecule resolution.

## INTRODUCTION

Parkinson’s disease (PD) is the second most prevalent neurodegenerative disorder after Alzheimer’s disease, affecting up to 3% of individuals over the age of 65^1^. A hallmark of PD is the progressive and irreversible degeneration of neurons, which correlates with characteristic motor symptoms. Post-mortem analyses of PD patient brains have consistently revealed the presence of intracellular protein aggregates, known as Lewy bodies, which are primarily composed of filamentous assemblies of *α*-synuclein (*α*S) proteins^2–5^. *α*S is a 140-residue intrinsically disordered protein (IDP) that is soluble under physiological conditions. It exhibits low hydrophobicity and a high net charge, and shares several aggregation-prone features with other amyloidogenic IDPs, such as A*β* in Alzheimer’s disease^6^. From a structural point of view, *α*S is often described as a “chameleon” protein^7–10^, remaining largely disordered in solution but capable of adopting local *α*-helical or *β*-sheet conformations under specific conditions, such as membrane binding or during oligomer formation^11–15^. Familial forms of PD have been associated either with overexpression of wild-type (WT) *α*S, often due to gene triplication, or with single-point mutations in the *α*S gene^16–21^. These missense mutations, including A30P and E46K among others, alter the aggregation kinetics and structural properties of the protein^22^. For instance, the E46K variant aggregates more rapidly than the WT, whereas the A30P variant aggregates more slowly. Moreover, the mutants differ not only in their fibril formation but also in the nature of their early-stage oligomers. A30P, in particular, shows a higher propensity to form non-fibrillar aggregates^19^. Given their distinct biochemical behaviors and pathological relevance, the propensity of sensors to detect and discriminate these variants at the single-molecule level holds significant promise for the development of early diagnostic biomarkers and a better understanding of PD-related neurodegeneration.

In this context, single-molecule technologies have gained increasing attraction in biotechnology research due to: i) their ability to capture the complex behaviour of biological molecules, and, ii) their unique capacity to probe molecular structure, dynamics and function, unhindered by the averaging inherent to ensemble experiments^23^. Among these approaches, solid-state nanopores (SSNs) represent a powerful, label-free platform for detecting and characterizing individual biological molecules^24–26^, from viruses, vesicles to proteins. The principle of detection relies on the resistive pulse technique: an SSN, typically a nanometer-scale hole drilled into a thin membrane, is immersed in an electrolyte solution, and a constant voltage is applied across the membrane. As a molecule translocates through the nanopore, it partially blocks the ionic current, producing transient current drops whose amplitude, duration, and shape serve as molecular fingerprints. Two-dimensional (2D) materials with nanometer-thick membranes are particularly well-suited due to their atomic-scale thickness, which enhances spatial resolution and sensitivity^27^. Among these, molybdenum disulfide (MoS_2_) has emerged as a promising candidate, offering larger signal-to-noise performance compared to other 2D materials such as graphene or boron nitride^28^. In addition to its favorable electrical, mechanical, and chemical properties^29^, MoS_2_ enables relatively precise detection of sequence motifs along translocating molecules^30–37^. For example, using bias MD simulations and machine learning techniques, it has been shown that single-layer MoS_2_ nanopores (*D* ≃ 1.85 nm) were able to detect individual amino acids in a polypeptide chain (16 units) with a high accuracy and distinguishability^31^. Moreover, we have recently demonstrated using unbias MD simulations that MoS_2_ nanopores (*D* ≃ 1.3 nm) can distinguish individual amino acids based on their charge, enabling coarse-grained sequencing of proteins^38^. Concurrently, experimental studies have shown that MoS_2_ nanopores (*D* ≃ 2.5 nm) were sensitive to polylysine translocation both in ionic and transverse signals^32^ and that MoS_2_ nanopores can resolve single amino acids with sub-Dalton precision^36^, discriminating chemical group differences and even identifying amino acid isomers (*D* ≃ 0.6 − 1.6 nm). Despite these promising developments, achieving accurate and robust protein sequencing with 2D nanopores remains a remarkable challenge, primarily due to the high translocation speed that restricts temporal resolution and hampers the identification of individual sequence features^39^.

Over the past decade, several experimental studies have been dedicated to the characterization of *α*S oligomers, which are increasingly recognized as the most cytotoxic species in the aggregation process, rather than mature fibrils^21,40,41^. Due to their structural heterogeneity, low concentration, and transient nature, probing these oligomeric intermediates represent a significant challenge to conventional biochemical techniques. Therefore, a diverse palette of single-molecule approaches has been applied to probe oligomer formation and dynamics with high sensitivity and resolution. It includes polymer-coated solid-state nanopores^42^, nanopipettes^43^, biological nanopores^44^, single-molecule fluorescence^45–49^, confocal spectroscopy^50,51^, two-color aggregate pulldown assays^52^, single-molecule force spectroscopy and optical tweezers^53–55^, as well as cryo-electron microscopy^56–59^. Collectively, these methods have significantly expanded our understanding of early *α*S aggregation events and have provided molecular insights into the pathways leading to neurotoxic oligomers. Nevertheless, while oligomeric states of *α*S have been extensively studied, the direct detection and sequencing of its monomeric form at the single-molecule level remains unexplored. In this context, 2D SSNs represent a powerful complementary approach. Rather than inferring protein states indirectly, nanopore sequencing enables the direct, label-free readout of individual amino acid sequences, capturing sequence variations, including point mutations implicated in familial PD. This molecular-scale resolution will open new possibilities for quantitative single-molecule diagnostics.

In this work, we performed all-atom bias molecular dynamics (MD) simulations in explicit solvent to investigate the full translocation of *α*S through MoS_2_ nanopores. This work represents, to the best of our knowledge, the first computational demonstration of nanopore sequencing of a protein biomarker with such length and structural disorder at the microsecond timescale, and characterized by ionic current traces resolved at near single-residue level. Explicitly, we studied the passage of the WT *α*S as well as two pathogenic variants, A30P and E46K. We performed five independent reads per variant, yielding to a total translocation time exceeding 22 *µ*s. For each read, we predicted ionic current traces analogous to those recorded experimentally, and leveraged the main advantage of MD, *i*.*e*. the ability to directly access atomic-level structural information throughout the translocation process. It allows us to establish the nonlinear relationship between amino acid position,orientation inside the pore, and the corresponding ionic current fluctuations. Our results provide new insights into the coupling between protein sequence, local conformation, and nanopore signal features. It represents a step toward nanopore-based protein sequencing and single-molecule biomarker identification. These findings lay the groundwork for future experimental validation using MoS_2_ SSNs.

## RESULTS AND DISCUSSION

### Amino Acid Sequence of Human *α*-synuclein Intrinsically Disordered Protein

*Human α*S is an IDP composed of 140 amino acids (14.46 kDa, UniProt P37840). Its primary structure, *i*.*e*. its sequence, is comprised of 17 out of the 20 natural amino acids (Fig. 1a), with arginine (R), tryptophan (W), and cysteine (C) completely absent. The most abundant amino acids in the sequence of *α*S are the hydrophobic valine (V) and alanine (A), present 19 times each, as well as glycine (G, 18), negatively charged glutamic acid (E, 18), and positively charged lysine (K, 15). Functionally, the sequence of *α*S can be divided into three regions with distinct biochemical properties (Fig. 1b): i) the N-terminal domain (N-term, residues 1–60), which is amphipathic and contains the majority of well-known missense mutations; ii) the NAC domain (residues 61–95), a highly hydrophobic region critical for fibril formation and aggregation; iii) the C-terminal domain (C-term, residues 96–140), which is highly acidic and contains several known phosphorylation sites.

**Figure 1.**
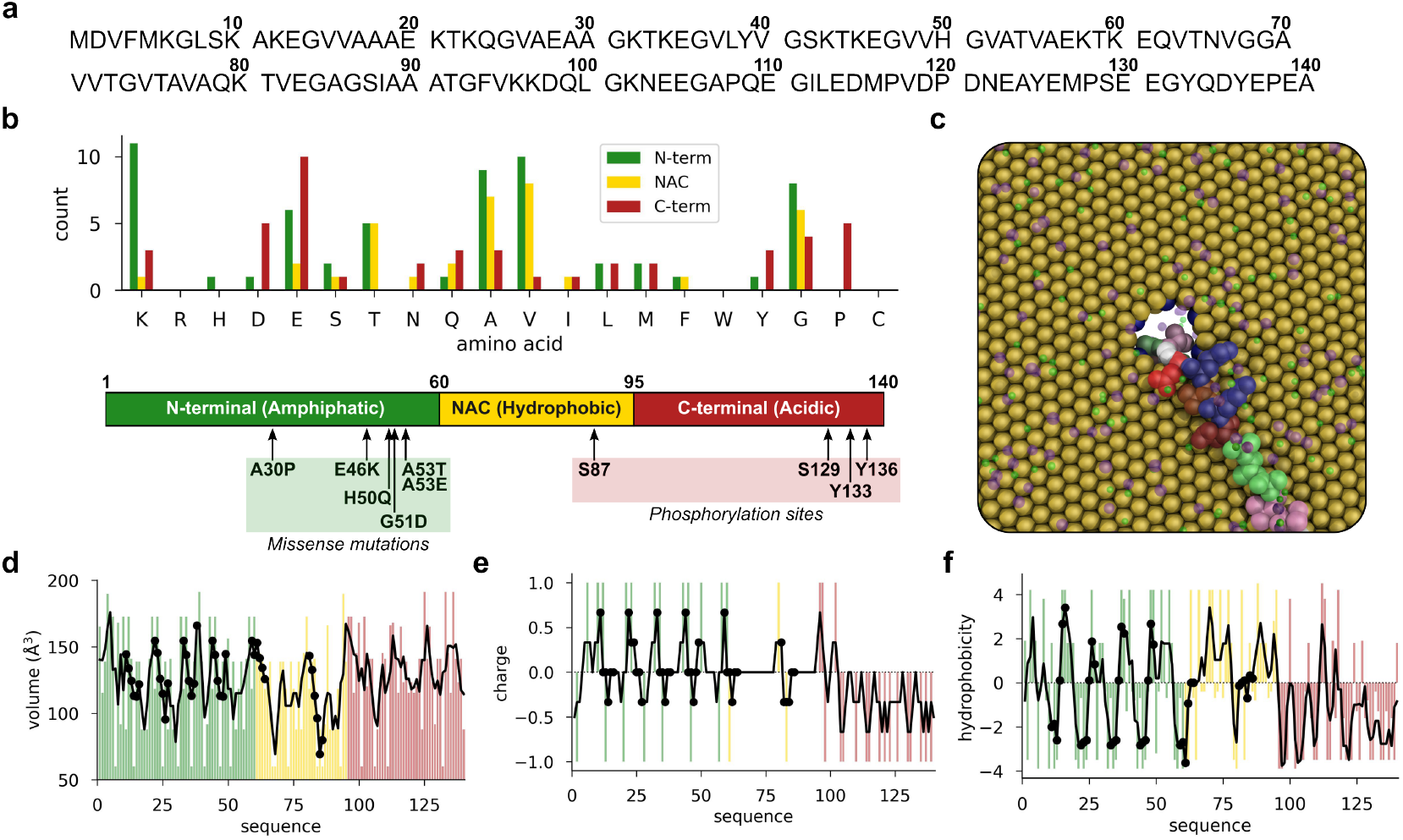
Amino Acid Sequence of Human *α*S Intrinsically Disordered Protein. a) Primary structure (sequence). b) Amino acid composition of the sequence of *α*S and its structural domain characteristics. The color code is the following: N-term (green), NAC (yellow) and C-term (red). c) Atomistic model of single-layer MoS_2_ SSN of diameter *D* ≃ 1.1 nm. d) Volume of amino acids along the sequence of *α*S. Black line indicates the moving average calculated using a window of 3 residues and black circles indicate the position of KTKEGV repeat motifs in the sequence (also shown in panels e and f). e) Net charge of amino acids along the sequence of *α*S. f) Hydrophobicity of amino acids along the sequence of *α*S, according to the Kyte-Doolitle scale^60^.

Several pathogenic mutations in the gene of *α*S, most notably single amino acid substitutions in the N-term domain (H50Q, G51D, A53E, A30P, A53T, and E46K), have been studied. They also have been linked to familial forms of PD and are of particular interest as biomarker. Moreover, the sequence of *α*S features six conserved repeat motifs (consensus: KTKEGV), which are also found shared in other members of the synuclein family (*β* - and *γ*-synuclein). These specific motifs are located in the sequence (Fig. 1a) at positions 10–15 (KAKEGV), 21–26 (KTKQGV), 32–37 (KTKEGV), 43–48 (KTKEGV), 58–63 (KTKEQV), and 80–56 (KTVEGA) and play a central role in oligomer formation, especially when missense mutations occur within these repeats^61^. Finally, as shown in Fig. 1d-f, these conserved motifs correspond to regions of reduced side-chain volume and charge, making them particularly important for nanopore-based sequencing approaches, which are highly sensitive to both properties^36,38,62^.

### Ionic Current Traces from Nanopore Sequencing of Wild-Type *α*-synuclein and Mutants A30P and E46K

Figure 2 shows the raw ionic current traces recorded during MD simulations of the full translocation of the WT *α*S and its pathogenic variants A30P and E46K through MoS_2_ nanopores. Panel a displays five independent reads for each protein sequence. Upon protein threading into the nanopore (see Movies S1 and S2), the ionic current rapidly drops, *i*.*e*. within a few tens of nanoseconds, from the open pore baseline of ≃ 1.6 nA (conductance of *D* ≃ 1.1 nm MoS_2_ nanopore) to near-zero nA, indicating maximum blockade (100%). During translocation, the traces exhibit pronounced fluctuations and multiple discrete levels, reflecting transient interactions between the amino acids and the pore. If, for most of the cases, amino acids translocate through the pore sequentially, ensuring an ordered passage, a very few number of “knots” were observed, corresponding to the passage of amino acids that are distant in the sequence by more than 3 amino acids (see Movie S1 with K_10_ for instance). During the passage of the *α*S protein, the ionic current varies between 0 and 1.0 nA, corresponding to a relative amplitude blockade of 40-100%, which is sufficiently broad to potentially discriminate specific sequence motifs within the sequence of *α*S protein. To evaluate the temporal resolution required to capture those features, we analyzed the information content of each ionic current trace. Sampling was performed every 10 ps over the course of microsecond-timescale MD trajectories. Using Shannon entropy as a metric (Fig. 2b), we found that the information content remains nearly unchanged (relative entropy close to 100%) up to a sampling interval of 1 ns. Beyond this threshold, the entropy starts to decrease, dropping to approximately 90% at a sampling interval of 100 ns. These results indicate that a time resolution of 1 ns is sufficient to resolve key ionic current fluctuations during *α*S protein translocation through MoS_2_ nanopores in these conditions.

**Figure 2.**
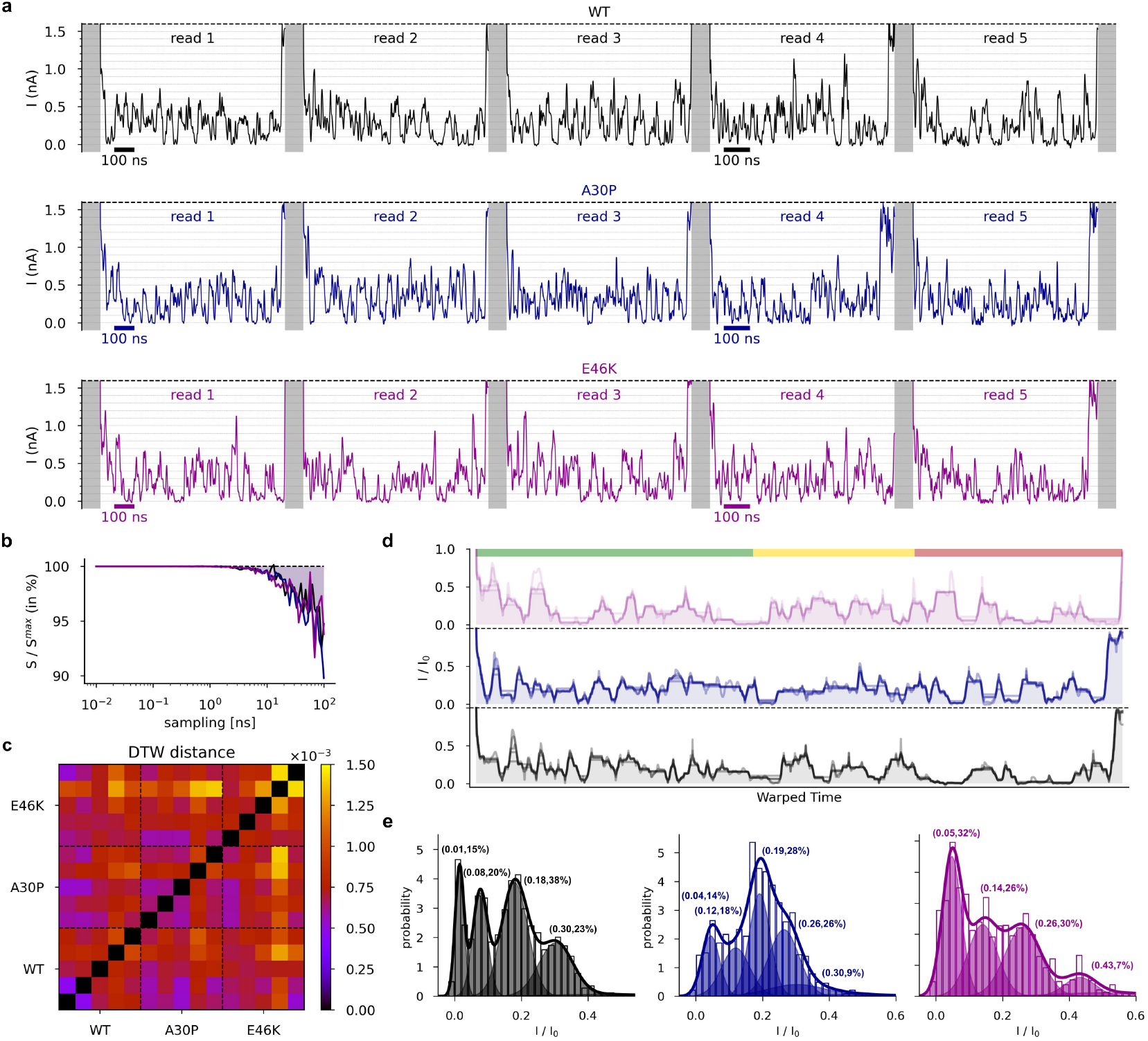
Ionic Current Traces from Nanopore Sequencing of Wild-Type *α*S and Mutants A30P and E46K. a) Ionic current traces recorded during MD production runs of the translocation of the WT *α*S (top panel, black), and its A30P (middle, blue) and E46K (bottom, magenta) mutants through a single-layer MoS2 nanopores. b) Relative Shannon entropy *S*/*S*^max^ as a function of the sampling interval (in ns), showing information content retained at different temporal resolutions. c) Distance matrix of DTW distances computed between all reads shown in panel a, comparing intra- and inter-sequence similarity. d) Ensemble ionic current traces (relative blockade current *I*/*I*_0_) obtained from aligned reads for the WT (black), A30P (blue), and E46K (magenta) *α*S. The color code is the same as in Fig. 1. e) Distribution of relative ionic current values derived from the clustering analysis of the ensemble traces shown in panel d, highlighting differences in blockade depth between WT and mutants.

We further compared the ionic current traces across multiple reads of the same sequence and between WT and mutant forms of *α*S. To do so, we performed dynamic time warping analysis (DTW, see Methods) to quantify the similarity between reads (Fig. 2c). This is particularly adapted for time series which may vary in speed. In the WT protein, the DTW distances among the five reads were relatively small, with reads 1 and 2 showing the highest similarity. A similar pattern was observed in the A30P mutant. In contrast, the E46K mutant displayed the largest variability, particularly between reads 3 and 5. When comparing the DTW inter-distances across WT, A30P, and E46K, the same trend holds, *i*.*e*. E46K exhibits the largest variations, especially between read 4 and all reads from WT and A30P *α*S. From the DTW distance matrix (Fig. 2c), we generated ensemble current profiles for WT, A30P, and E46K by aligning individual traces (Fig. 2d). While these ensemble traces preserve the multilevel features observed in the raw data (Fig. 2a), several characteristics emerge. Notably, WT and E46K both display deeper and more sustained current blockades than A30P, with overlapping features particularly evident in the transition from the N-term region to the NAC domain. In contrast, this transition appears shorter and less pronounced in A30P. E46K also presents extended blockade events in the N-term region, a domain where WT and A30P show only moderate fluctuations. In the C-term region, the WT trace shows longer and more pronounced current drops compared to both mutants. These differences shows how single-point mutations within the N-term domain can significantly alter the ionic current profiles during translocation through MoS_2_ nanopores.

Furthermore, we performed a clustering analysis on the ensemble traces for each *α*S variant to quantify how single-point mutations affect the distribution of relative current blockade. As shown in Fig. 2e, distinct patterns are observed across the different proteins. First, for the WT *α*S, six clusters were identified: four dominant ones within the 0-0.6 *I*/*I*_0_ range, each contributing over 15% of the total weight; and two minor clusters beyond 0.6 *I*/*I*_0_ (< 40% blockade), each accounting for less than 2%. The most prominent cluster (38% weight) is centered at 0.18, corresponding to 82% blockade, followed by a second cluster (23%) centered at 0.30 (70% blockade). Two additional clusters, located in the deep blockade regime at 0.08 (92%) and 0.01 (99%), contribute to 15% and 20% respectively, for a total of 35% weight. Compared to the WT *α*S, A30P presents a total of seven clusters, including five dominant ones and two with minor contributions. In addition, A30P shares a similar most populated cluster centered around 0.19 (28% weight), followed by a characteristic second cluster (26%) centered around 0.26. A smaller cluster at moderate blockade (0.30, 9%) is also observed. However, compared to WT, A30P shows a reduced contribution from deep blockade events: the most pronounced cluster in this regime is centered at 0.04 (14% weight), while another at 0.12 accounted for 18%. These differences further support the attenuated current variations observed in the raw data for A30P (Fig. 2a). In contrast, the E46K mutant exhibits a remarkably different distribution (Fig. 2e), dominated by deeper current blockades. Its most populated cluster (32%) is centered at 0.05 (95% blockade), accompanied by two additional clusters of comparable weight, centered at 0.14 (26%) and 0.26 (30%). Compared to the WT and the A30P mutant, E46K shows a distinct shift toward deeper blockade levels and a notable reduction in moderate blockades. Overall, the WT *α*S trace distribution appears as a hybrid of the two mutants, encompassing both moderate and pronounced blockades. A30P exhibits a tendency toward weaker blockade patterns, while E46K favors stronger ones. This population shift in E46K clustering analysis likely results from the substitution of a negatively charged glutamic acid with a positively charged lysine in the N-term domain, enhancing electrostatic interactions within the local electric potential inside the nanopore and increasing current fluctuations during translocation. While ensemble traces and blockade distributions from clustering analysis provide valuable global comparisons between the WT and mutant *α*S proteins, a deeper investigation is necessary to interpret transient events and discrete transitions in ionic current traces. To this end, we extracted the stepwise signals from the raw traces shown in Fig. 2a to better characterize individual blockade events and their sequence-dependent properties.

### Stepwise Signals of Ionic Current Traces for Wild-Type *α*-synuclein and Mutants A30P and E46K

Figure 3 shows a representative stepwise signal (WT read 1, see also Movies S1 and S2), obtained from the raw ionic current trace presented in Fig. 2a. Additional signals of the 15 reads are provided in the Supplementary Information (Fig. S1). First, the Pearson correlation between the raw and stepwise signals is consistently very high across all reads, indicating that the stepwise transformation accurately preserves the overall signal features. Correlation values range from 0.979 (min, E46K read 4) to 0.988 (max, E46K read 5). Second, the average number of steps detected per translocation event is approximately 55 *±* 5 for the WT *α*S protein and 57 *±* 7 for both A30P and E46K mutants (Fig. 3b). Compared to the full-length sequence of *α*S (Fig. 1), this corresponds to roughly 40%, which implies that individual amino acids are not totally resolved. Instead, each step reflects the passage of short sequence motifs, usually consisting of 1 to 3 amino acids, though occasionally involving longer segments (less frequent). This limited resolution can be attributed to two main factors: i) heterogeneous translocation speeds between different amino acids, and ii) sequence-dependent effects, such as side-chain volume, net charge, or hydrophobicity (Fig. 1d–f), that influence both the duration *τ* (dwell time) and relative depth *I*/*I*_0_ of the current blockade.

**Figure 3.**
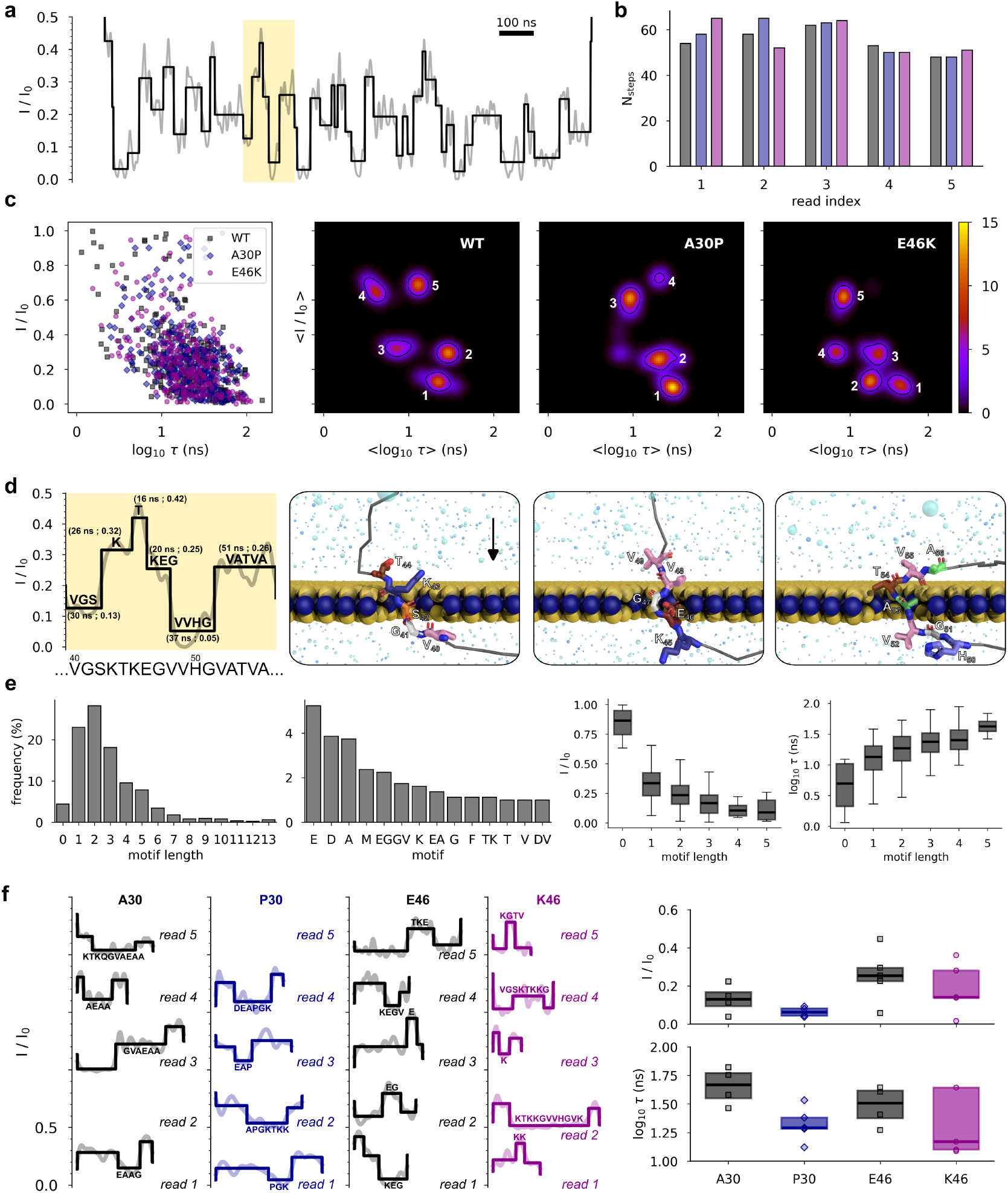
Stepwise Signals of Ionic Current Traces for Wild-Type *α*S and Mutants A30P and E46K. a) Representative stepwise signal derived from the ionic current trace (WT, read 1). b) Number of discrete steps (blockade events) detected per read for each protein variant. c) Clustering analysis of step characteristics based on dwell time (*τ*, in ns) and relative current blockade (*I*/*I*_0_). *I*/*I*_0_ and *τ* in brackets indicate cluster means obtained from BGM. d) Correlation between individual blockade steps and the underlying sequence motifs. The trace corresponds to the yellow-highlighted segment in panel a. Atomic representations illustrate the translocation of identified sequence motifs through the pore, as observed in MD simulations. The direction of translocation (indicated by the black arrow) is from the top of the membrane to the bottom, identical to the direction of the electric field. e) Statistical distribution of sequence motifs detected across all reads. Histograms (left panels) represent the frequency (in %) of motif length and sequence. f) Blockade events associated with mutation at position 30 and 46. Comparative boxplots of dwell time and relative current blockade are shown for WT, A30P, and E46K reads.

Figure 3c summarizes the distributions of these parameters; detailed statistics for each sequencing datasets are available in the Supplementary Information (Fig. S2). First, dwell time *τ* ranges from a few nanoseconds to several hundred of nanoseconds, with the most frequent values exceeding several tens of nanoseconds. This is consistent with previously reported timescale for peptide translocation through single-layer MoS_2_ nanopores^33,34,37,38^. Moreover, relative blockade values span from 0 (complete current blockade) to 1 (fully open pore), with most events falling between 0 and 0.4, indicating substantial steric and/or electrostatic interactions within the nanopore. Interestingly, a strong correlation is observed between dwell time and blockade amplitude, *i*.*e*. longer events tend to correspond to deeper blockades, suggesting stronger interactions with the pore. Conversely, short translocation events (*e*.*g*., *τ* < 10 ns) typically show minimal blockade (*I*/*I*_0_ > 0.8), which may reflect either weak interactions within the pore or the rapid passage of small, disordered peptide fragments.

Based on the blockade characteristics described above, we performed clustering analysis to identify and group together the most relevant signal fingerprints within the stepwise ionic current traces of WT *α*S and its A30P and E46K mutants (see Methods). As shown in Fig. 3c (see also Table 1), the WT protein is characterized by five distinct clusters. Among these, three clusters are characterized by moderate to deep relative blockade values (*I*/*I*_0_ < 0.35) and intermediate dwell time, ranging from 8 to 30 ns. The two remaining clusters exhibit smaller blockade levels (centered around *I*/*I*_0_ ≈ 0.65) and shorter dwell time, between 4 and 13 ns. The most prominent cluster (index 1), centered at (22 ns; 0.13, 87% blockade), accounts for approximately 49% of the total signal weight, followed by cluster 2 (29 ns; 0.29, 0.71) with 30% (Tab. 1). The remaining three clusters contribute marginally, with weights ranging from 5 to 9%. In comparison, the A30P mutant is characterized by four clusters. Clusters 1 and 2 are highly consistent with those of the WT, together encompassing nearly 80% of the total signal. However, cluster 3, which is present in WT with non-negligible weight, is absent in A30P (weight < 1%), suggesting a mutation-induced loss of that specific signal fingerprint. The two clusters corresponding to smaller blockade levels (*I*/*I*_0_ > 0.6, blockade < 40%) are also present in A30P and remain qualitatively similar to those in the WT, though they exhibit slightly prolonged dwell time in the mutant (5-20 ns). In addition, the E46K mutant presents five clusters. Four of them are located in the deep blockade regime (*I*/*I*_0_ < 0.3, blockade > 70%), as opposed to three in the WT, while only one cluster falls in the high-blockade regime (compared to two in the WT). Remarkably, cluster 1 in E46K is centered at (42 ns; 0.11), corresponding to the longest dwell time observed among all datasets. This suggests that the E46K mutation induces the translocation of longer sequence motifs through the nanopore, possibly due to altered electrostatic or conformational properties of the local sequence environment. This interpretation is consistent with the known biochemical effect of the E46K substitution, which replaces a negatively charged glutamic acid with a positively charged lysine, thereby drastically altering the local and global charge decoration of *α*S. These results align with previous findings indicating that MoS_2_ nanopores are highly sensitive to charge and can discriminate between negatively and positively charged amino acids based on their characteristic blockade profiles^38^.

**Table 1.** Clustering analysis of stepwise ionic current signals recorded during the sequencing of WT *α*S and its A30P and E46K mutants through MoS_2_ nanopores. For each cluster index, the table lists: (i) the number of data points assigned to the cluster, (ii) the cluster’s relative weight (percentage of total signal), and (iii) the cluster centroid coordinates, given as (dwell time *τ* in ns; relative blockade *I*/*I*_0_)

| Cluster index | Wild-Type | mutant A30P | mutant E46K |
| --- | --- | --- | --- |
| 1 | 844 / 4510<br>(22 ns ; 0.13)<br>49% | 935 / 3129<br>(28 ns ; 0.11)<br>35% | 829 / 4091<br>(42 ns ; 0.12)<br>16% |
| 2 | 978 / 4510<br>(29 ns ; 0.29)<br>30% | 961 / 3129<br>(20 ns ; 0.26)<br>51% | 905 / 4091<br>(19 ns ; 0.14)<br>31% |
| 3 | 802 / 4510<br>(8 ns ; 0.32)<br>9% | 954 / 3129<br>(9 ns ; 0.60)<br>13% | 845 / 4091<br>(21 ns ; 0.29) |
| 4 | 914 / 4510<br>(4 ns ; 0.65)<br>5% | 279 / 3129<br>(20 ns ; 0.73)<br>1% | 601 / 4091<br>(7 ns ; 0.30)<br>6% |
| 5 | 972 / 4510<br>(13 ns ; 0.69)<br>7% | - | 911 / 4091<br>(8 ns ; 0.62)<br>12% |

Next, the relationship between individual blockade events in the stepwise ionic current signal and the underlying amino acid sequence was examined through the identification of sequence motifs associated with each discrete step (see Methods). A representative example, extracted from the WT read 1 (highlighted in yellow in Fig. 3a), is presented in Fig. 3d. This segment corresponds to the translocation of a sequence spanning residues V40 to A56, with a total duration of approximately 200 ns. Within this interval, six distinct motifs were identified: one composed of five amino acids, one of four, one of three, and two comprising single amino acid. Motifs consisting of 5 residues generally exhibited the longest dwell time (Fig. 3e), though they did not systematically correspond to the deepest current blockades, highlighting the multifactorial nature of ionic current signal fluctuations. On the other hand, single-residue motifs displayed the shallowest relative blockades, yet their dwell time could match those of longer motifs, suggesting transient interactions or delayed release events within the sensing region. Motifs of length zero correspond to open pore current levels, with no amino acid associated to them due to very fast translocation speed through the nanoporous membrane. They are typically observed at the initial entry and final exit of the protein in the pore. Aggregating the analysis across the 15 nanopore reads, we found that MoS_2_ single-layer nanopores predominantly resolve short motifs composed of one to three amino acids, consistent with previous reports^34^. Among the most frequently detected residues are glutamic and aspartic acid (D), both negatively charged, followed by small hydrophobic residues such as alanine, methionine (M), and valine. Glycine, due to its minimal side-chain volume, is rarely detected as an isolated event. Instead, it frequently appears in two-residue motifs, *e*.*g*. EG or GV, suggesting that its translocation through the pore does not produce a sufficiently strong blockade on its own, but becomes detectable when it co-translocates with a neighboring amino acid. These findings support the idea that detection is governed not only by molecular volume but also by electrostatic interactions, as evidenced by the pronounced blockade signatures associated with positively charged lysine, particularly in TK-containing motifs (threonine, lysine). The intrinsic amino acid composition of *α*S sequence, where E, D, K, A, V, and G are among the most recurrent residues, likely contributes to the higher probability of observing these residues in the stepwise signal profiles.

Finally, we extracted signal segments corresponding to the translocation of amino acids 30 and 46, which are the locations of the missense mutations under investigation. However, due to the intrinsic resolution limits of MoS_2_ nanopores which do not permit residue-level discrimination along the sequence from the N-to the C-termini, plus the stochastic nature of single-molecule translocation in SSNs, it is not feasible to unambiguously associate individual blockade events with specific amino acids at precise sequence positions. Instead, we analyze short sequence motifs encompassing the residue of interest and characterize their collective translocation signatures. For mutation site 30, the substitution of alanine with proline (P) does not substantially alter the relative blockade levels compared to dwell time, despite the known conformational rigidity introduced by proline. A charged motif, KTKQGV, located at positions 21–26, dominates the stepwise signal recorded during MD simulations (Fig. 3f), suggesting that local volume and charge variations (Fig. 1d-f) play a significant role in current modulation in this region. In contrast, for mutation site 46, the replacement of glutamic acid with lysine induces more pronounced signal variability. Dwell time in this region vary strongly, depending on the length of the motif being translocated (ranging from 1 to 11 amino acids). This mutation site is embedded within the KTKEGV motif (positions 43–48), and the mutation-induced shift in charge decoration strongly influences both the extent and duration of translocated motifs, thereby modifying the blockade pattern. These findings reinforce the sensitivity of 2D nanopores to electrostatic chemical environments of amino acids and charge distribution within protein sequences.

### Sequence Fingerprints from Ionic Current of Wild-Type *α*-synuclein and Mutants A30P and E46K

In the context of nanopore sensing, molecular dynamics operates as a computational nanoscope, providing atomistic-resolution insight into the conformational dynamics and molecular interactions occurring during protein translocation. MD trajectories, which comprise time-resolved Cartesian coordinates of all atoms in the system (membrane, protein, electrolyte) at the picosecond timescale, enable the reconstruction of the complex, non-linear relationship between the amino acid passage through the pore and the resulting ionic current fluctuations observed during nanopore sequencing. To elucidate this relationship, we quantified, at each timestep, the instantaneous volume of amino acids located within the nanopore sensing region. In contrast to the sequence motifs uniquely associated to stepwise signals as shown in Fig. 3, an ionic current value at a given time *t* can be associated to different amino acids. While such spatio-temporal resolution is inaccessible experimentally, this computational approach proves highly informative, offering direct access to the physical mechanisms underlying signal fluctuations in MoS_2_-based nanopore systems.

Figure 4a presents the amino acid volume inside the pore and the corresponding ionic current trace for the representative WT read 1 (see also Movies S1 and S2). The other 14 traces are shown in Supplementary Information (Fig. S3). First, the instantaneous volume fluctuates between 70 and 230 Å^3^, consistent with the known volume range of individual amino acids (*e*.*g*., from glycine ~ 60 Å^3^, to tryptophan ~ 228 Å^3^). This confirms that single-layer MoS_2_ with *D* ~ 1.1 nm pores offer suitable resolution to detect amino acid-level volume variations. Second, significant increases in volume trace occur at specific positions along the sequence of *α*S. For instance, the transition between the N-term and NAC domain is sharply defined in almost all reads, except WT read 4, A30P read 5, and E46K read 1. This transition coincides with the KTKEQV motif, featuring lysine residues with positively charged and relatively long side chains. A similar transition between the NAC and C-term domain is consistently observed, particularly in the sequencing of the WT protein compared to the mutants. Next, we evaluated the correlation between volume fluctuations and ionic current traces at different timescale. As shown in Fig. 4b, the total Pearson correlation coefficient is significantly negative across all protein variants: − 0.61 *±* 0.06 for WT, − 0.65 *±* 0.05 for A30P, and − 0.56 ± 0.13 for E46K. This indicates that increases in confined volume are generally associated with deeper ionic current blockades. The weaker and more variable correlation in E46K reflects the mutation-induced disruption of charge decoration in the sequence, which decreases the influence of volume on current fluctuations. In addition, dynamic correlation analysis along the traces, computed on sliding windows of 50 ns along the traces, confirms a predominantly anti-correlated behavior (*ρ* < 0). The distribution shown in Fig. 4b presents four distinct components: two major ones centered at *ρ* = − 0.74 and *ρ* = − 0.92, which account for 50% of the data, and coincide with strong current blockades. A third, less frequent component is centered around *ρ* = 0.31, with values reaching up to *ρ* = +1 in localized regions, suggesting that other factors, such as electrostatic interactions and local protein conformation, prevail over volume-related effects.

**Figure 4.**
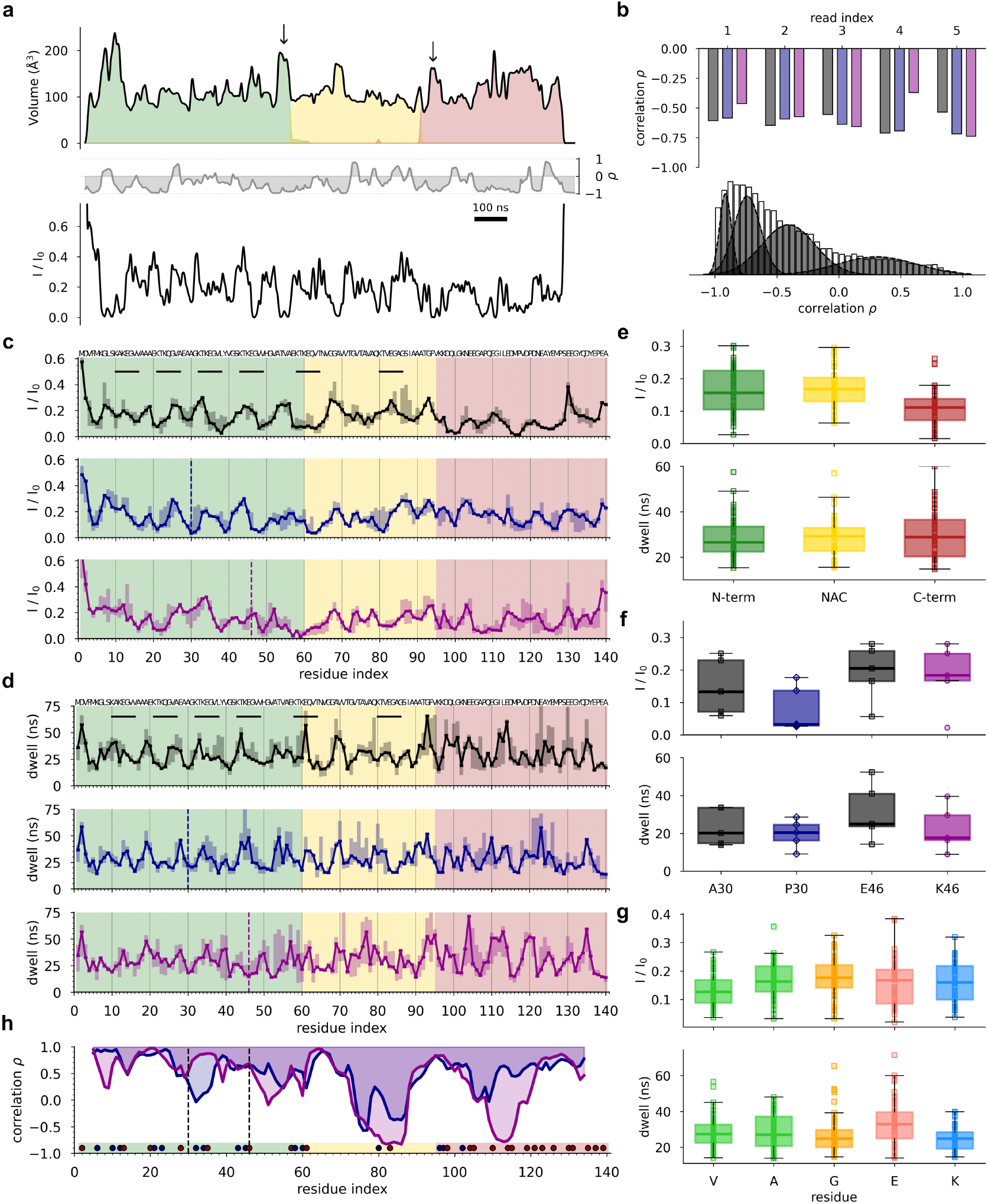
Sequence Fingerprints from Ionic Current of Wild-Type *α*S and Mutants A30P and E46K. a) Volume of amino acids inside the nanopore (in Å^3^) and the corresponding ionic current trace recorded during the translocation of WT *α*S (read 1). Middle panel shows the dynamic correlation *ρ* over time between the two traces. Black arrows show large volume changes between the different sequence domains. The color code is the same as in Fig. 1. b) Statistical analysis of correlations *ρ* across all reads. Top panel indicate the total correlation computed between the full traces shown in panel a and in Fig. S3. Bottom panel shows the distribution of dynamic correlation over time including datasets from the 15 reads. c) Ionic current profile along the sequence of *α*S. KTKEGV repeat motifs are indicated with horizontal bars. d) Dwell time profile (in ns) along the sequence of *α*S. e), f), g) Boxplots of ionic current and dwell time for protein domains, mutation sites, and most frequent residues, respectively, including datasets from the 15 reads. h) Correlation *ρ* as a function of residue index between current profiles shown in panel c for WT vs. A30P (blue) and WT vs. E46K (magenta). Mutations at position 30 and 46 are indicated with dashed lines; charged residues are also marked (blue: positive, red: negative).

Leveraging the volume trace extracted from MD, we assigned each ionic current value over time to the specific amino acids present within the sensing region. This assignment, only achievable in MD, enables the construction of sequence-specific fingerprints (*i*.*e*. profiles) of current blockades and dwell time for *α*S, as shown in Fig. 4c and d. First, these fingerprints show strong similarities across WT and mutant proteins. The Pearson correlation between ionic current profiles is 0.64 (WT vs. A30P) and 0.71 (WT vs. E46K), while for dwell time profiles, the correlation is 0.64 and 0.54, respectively. Median current blockade levels (excluding N- and C-termini) vary between 0 and 0.35. Interestingly, as observed from DTW analysis (Fig. 2) and stepwise analysis (Fig. 3), the E46K mutant exhibits deeper average blockades, while the A30P mutant tends to show shallower ones, particularly in the C-term domain (Fig. 4e). Distinct local patterns emerge along the sequence, often associated with clusters of charged residues: notably, the KTKEGV motifs in the N-term region and the poly-glutamate stretches in the C-term domain. Second, at the mutation sites, we observe differential effects. The A30P mutation leads to the largest localized change in the ionic current profile, whereas the E46K mutation induces the largest difference in dwell time (Fig. 4f). Second, we analyzed dwell time profiles along the sequence (Fig. 4 d). These range from 10 to 70 ns, with negatively charged residues (*e*.*g*., glutamic acid) showing longer residence times inside the pore, while positively charged residues (*e*.*g*., lysine) and glycine show shorter dwell times (Fig. 4 g).

These effects highlight how substitutions can strongly impact structural rigidity (proline) versus charge decoration (lysine), both of which modulate translocation dynamics. To localize sequence regions most affected by the mutations, we computed the moving average correlation (Fig. 4h) using a sliding window of size equal to 10 residues between the WT and mutant current profiles. We then plotted the results in shown in Fig. 4c. First, we observed that the A30P mutation alters current fluctuations near the mutation site (residues 28–36), but also affects distant regions, including residues 72–94 (in the NAC domain), which contains the final KTVEGA motif and a high density of small amino acids (glycine, alanine). Similarly, residues 104–112 in the C-term domain are impacted, likely due to local enrichment in negatively charged or small residues. Second, similar patterns are observed for the E46K mutant: in addition to strong alterations near the mutation site (residues 46–58), regions 72–94 and 104–122 are also significantly affected. While there is overlap in the regions impacted by both mutations, E46K appears to influence a broader segment of the C-term domain, likely due to its more substantial influence on charge distribution. Overall, these findings are consistent with previous studies^37,38^, and further support the interpretation that both electrostatics and molecular size contribute to translocation dynamics. Together, these results demonstrate that single-point mutations in *α*S can significantly affect ionic current signatures, not only at the site of mutation but also at distant sequence regions. This also supports the hypothesis that such mutations perturb long-range intra-molecular interactions and global folding propensities, rather than acting locally alone^10,63^. These insights underline the capacity of 2D nanopore-based sequencing, coupled with MD simulations, to resolve sequence-specific features of intrinsically disordered proteins at the nanoscale.

## CONCLUSION

In this study, we provide a comprehensive multi-scale investigation of ionic current signals recorded during the translocation of wild-type and mutant *α*-synuclein proteins through MoS_2_ nanopores. By integrating dynamic time warping analysis of raw current traces, stepwise ionic current segmentation, unsupervised learning clustering, and MD-informed (volume) sequence fingerprinting, we offer a detailed and mechanistic understanding of how such an intrinsically disordered protein sequence features and single-point mutations modulate nanopore variations across multiple temporal and spatial resolution. First, we showed that the WT *α*S exhibits a characteristic translocation signature with well-aligned ionic current traces across multiple reads, indicating reproducible global dynamics. Stepwise signal segmentation reveals discrete blockade events corresponding to the passage of short sequence motifs, typically one to three amino acids in length. Statistical clustering of these steps identifies distinct signal populations differing by dwell time and blockade amplitude, which are correlated with local physico-chemical properties of the translocating residues, particularly volume and charge. MD simulations further elucidate the underlying physical basis, confirming that ionic current fluctuations strongly anti-correlate with the instantaneous volume of amino acids inside the nanopore, while deviations highlight the role of electrostatic interactions and local conformational dynamics. Sequence fingerprinting based on MD trajectories associates specific blockade and dwell time patterns with sequence motifs, including charged residues that particularly influence current modulation.

Second, we investigate the impact of two PD-associated point mutations, A30P and E46K. Both mutations produce well-marked alterations in the ionic current signatures, but through distinct mechanisms. The A30P mutation, which introduces a conformationally rigid proline residue, leads to the disappearance of certain clusters of blockade events and modifies the step distribution, particularly affecting regions distant from the mutation site, such as in the NAC and C-terminal domains. This suggests that A30P perturbs the overall protein dynamics and structural flexibility during translocation. In contrast, the E46K mutation, which replaces a negatively charged glutamic acid with a positively charged lysine, induces levels and bumps with longer dwell time and altered blockade amplitudes within charge-sensitive motifs, reflecting substantial local electrostatic effects. Finally, dynamic correlation analyses along the sequence demonstrate that both mutations not only impact the immediate sequence neighborhood but also modulate ionic current features in distant protein regions. This is consistent with long-range intramolecular interactions characteristic of intrinsically disordered proteins^10,63^.

Overall, our approach combining atomistic MD simulations and nanopore datasets (ionic current) analysis highlights the sensitivity of MoS_2_ nanopores to sense molecular variations in protein sequence and local conformation. The ability to resolve distinct sequence motifs and detect mutation-induced changes in ionic current patterns paves the way toward nanopore-based single-molecule protein sequencing for early disease diagnostics applications. These findings emphasize the promise of two-dimensional material nanopores as versatile platforms for probing complex biomolecular dynamics with extraordinary resolution.

## METHODS

### Atomistic Modeling of MoS_2_ Solid-State Nanopores for Sequencing Applications

The SSN system modeled in this study consists of a monolayer MoS_2_ membrane and one *α*S protein (wild-type or mutant), both immersed in a 1 M KCl electrolyte solution. To simulate the complete translocation process, the protein was initially positioned approximately 2.5 nm above the nanopore. The full atomistic system includes ≃ 540, 000 atoms, encompassing the MoS_2_ membrane, the protein, water molecules, and ions. The MoS_2_ membrane was constructed using an orthorhombic unit cell of 2H-MoS_2_, with lattice vectors 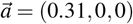 and 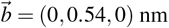, containing six atoms (two Mo and four S). The Mo–S bond length was set to 0.24 nm, and the interlayer S–S spacing to 0.32 nm, corresponding to the geometrical thickness of the membrane; the effective thickness being approximately 0.96 nm^64^. A cylindrical pore was drilled at the center of the membrane by removing atoms whose positions satisfy *x*^2^ + *y*^2^ < *R*^2^, yielding a pore diameter *D* ≃ 1.1 nm. The lateral dimensions of the membrane are 25 *×* 25 nm^2^. To model electrostatics accurately, atomic partial charges for Mo and S atoms were assigned using charge equilibration in vacuum based on ReaxFF^65^, ensuring overall charge neutrality of the membrane. This approach captures the influence of pore surface charges on ionic conductance with greater physical fidelity^66,67^. Initial atomic structures of *α*S (WT, A30P, and E46K) were generated from their amino acid sequences using AmberTools^68^. All proteins were modeled in a fully unfolded state using the LEaP module. Finally, explicit water using TIP3P model^69^, K^+^, and Cl^−^ ions^70^ were added to the system at 1 M concentration using the solvate module of GROMACS^71^.

### Molecular Dynamics Simulations of Protein Translocation

We performed all-atom classical molecular dynamics in explicit solvent to simulate the translocation of the intrinsically disordered protein *α*S (WT and mutants A30P, E46K) through MoS_2_ nanopore. MD simulations were carried out using the GROMACS 2023.3 software package^71^, with the AMBER99SB*-ILDN-q force field^72,73^for the protein. Force-field parameters for MoS_2_ are given in details in previous works^37,38^. Neighbor searching was performed by using a pair list generated using the Verlet method (particle-based cutoffs) as implemented in GROMACS^71^. The neighbor list was updated every 10 fs, with a cut-off distance for the short-range neighbor list of 1.0 nm. Moreover, electrostatic interactions were computed using a Coulomb potential and Van der Waals interactions using a Lennard-Jones (LJ) potential plus arithmetic mixing rules. Technically, Particle-Particle Particle-Mesh (PPPM) method^74^ was used to describe long-range electrostatic interactions with a Fourier spacing of 0.16 nm and a PME order of 4. A cutoff of 1.0 nm was applied to both Coulomb and LJ potential for non-bonded interactions. Finally, a long-range analytical dispersion correction was applied to the energy and pressure.

Prior to production, each system was equilibrated in two stages: first under constant volume and temperature (NVT, *T* = 300 K) and then under constant pressure and temperature (NPT, *P* = 1 bar), both for 100 ps, and without any applied electric field. These equilibration steps, performed using a time step of 1 fs and position restraints applied to the protein, allowed the system to stabilize thermally and structurally. Then, production runs were initiated by assigning random initial velocities and applying a uniform external electric field of 0.1 V/nm along the *z*-axis, corresponding to a transmembrane bias of 1 V. To promote translocation, pulling with umbrella potential was applied between the N-terminal methionine amino acid of the protein and the center of the nanopore, using cylindrical coordinates (*ρ, z*). For the normal coordinate *z*, two different pulling rates were used: 3.10^−5^ nm/ps for readouts 1–3 and 5.10^−5^ nm/ps for readouts 4–5 (with a force constant of 1000 kJ/mol/nm^2^), resulting in different trajectory duration, ranging from 1.02 *µ*s (reads 4–5) to 1.78 *µ*s (reads 1–3). For the radial coordinate *ρ*, pulling rate was set to zero (with a force constant of 600 kJ/mol/nm^2^). Production simulations were run in the NVT ensemble at 300 K using a time step of 2 fs with bonds involving H atoms constrained using the LINCS algorithm^75^. In total, five independent translocation simulations were performed for each protein sequence (WT, A30P, E46K), leading to 15 MD trajectories and a cumulative simulation time of 22.1 *µ*s. These simulations required over two million CPU hours at MesoBFC (see Acknowledgments), with an average scaling of ≃ 400 ns/day on 1392 cores. Altogether, MD simulations generated approximately 2.6 TeraBytes of translocation data.

### Ionic Current Traces recorded during Molecular Dynamics

Ionic current traces due to blockade events observed during the translocation of *α*S through MoS_2_ nanopores were computed from MD production runs using trajectories of K^+^ cations and Cl^−^ anions *i* over time *t*, which align with the electric field, *i*.*e. z*_*i*_(*t*), given the equation:

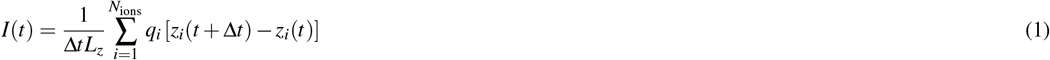

where Δ*t* represents the time interval between MD snapshots extracted for the calculations (Δ*t* = 1 ns); *L*_*z*_ corresponds to the dimension of the simulation box in the *z*-direction (*L*_*z*_ = 10 nm), which aligns with the applied electric field direction; *N*_ions_ corresponds to the total number of ions in the simulation box (*N*_*ions*_ = *X*), *q*_*i*_ is the charge of ion *i* (*q*_*i*_ = *±* 1), and *z*_*i*_(*t*) corresponds to the *z*-coordinate of ion *i* at time *t*.

Ionic current traces were computed every 10 ps along MD production runs and traces were filtered to remove high-frequency fluctuations by applying a low-pass Bessel filter with *N* = 4 and *W*_*n*_ = 0.03 (reads 1-3) and *W*_*n*_ = 0.02 (reads 4-5) using the `scipy` Python library (version 1.14.0). We also defined the relative ionic current as the ratio between the blockade current *I* and the open pore current *I*_0_. Mean and standard deviation values of open pore current of a single-layer MoS_2_ nanopore of diameter 1.1 nm are 1.60 and 0.15 nA and were extracted from the I-V curve (in the range 0-1 V) generated from independent MD simulations^76^ (similar MD parameters but no protein in the simulation box). Using this parameter, a ratio of zero indicates 100% blockade and a ratio of 1 a blockade of 0%.

#### Dynamic Time Warping Analysis

To quantify the similarity between ionic current traces, we computed the Dynamic Time Warping (DTW) distance between all pairs of traces, as implemented in the `dtai distance` Python library (version 2.3.12)^77^. DTW distances were normalized by dividing them by the sum of the lengths of the two signals, to account for variability in translocation duration. To generate ensemble traces for each *α*S sequence, we first identified the trace with the lowest mean DTW distance to all other traces and stretched it to create the so called medoid trace, **T**_**m**_ = [*t*_1_, *t*_2_, …, *t*_*n*_], where *n* is the mean length of all traces. We then DTW-aligned every other trace to **T**_**m**_ and created the so called consensus trace as the median of all aligned traces, **T**_**c**_ = [median(alignments to *t*_1_), median(alignments to *t*_2_), …, median(alignments to *t*_*n*_)]. Finally, ensemble traces in Fig. 2d show all traces aligned to the **T**_**c**_, but do not show **T**_**c**_. This method was adapted from a very recent work about multi-pass, single-molecule nanopore reading of long protein strands^62^.

#### Current Trace Step Analysis

To identify discrete ionic current levels during *α*S translocation through MoS_2_ nanopores, traces were segmented using the Pruned Exact Linear Time (PELT) algorithm^78^ implemented in the `ruptures` Python library (version 1.1.9). This method detects abrupt changes, *i*.*e*. changepoints, in the signal by minimizing a cost function based on the sum of squared deviations within each segment (L2 model). In this work, each current trace was treated as a one-dimensional time series, and the PELT algorithm was applied with a fixed penalty value to control the number of detected steps. The choice of penalty was empirically determined to balance the sensitivity to small fluctuations and the avoidance of over-segmentation. The detected changepoints correspond to transitions between relatively stable current levels, and each segment between two changepoints was considered a step. For each step, we extracted two main features: i) its dwell time *τ*, corresponding to the duration of the segment and representing how long a sequence motif remains in the sensing region of the pore; ii) its relative current blockade *I*/*I*_0_, which corresponds to the mean blockade current within the segment, normalized by the open pore current. This approach enabled quantitative characterization of translocation events at sub-sequence resolution and facilitated comparison across WT and mutant *α*S proteins.

#### Gaussian Mixture Clustering Analysis

Clustering of 1D and 2D datasets was performed using a Bayesian Gaussian Mixture (BGM) model, as implemented in the `scikit-learn` library (version 1.5.1). The optimal number of components for each dataset, denoted *N*_*c*_, was defined as the number of components with a weight greater than 1%. For each value of *N*_*c*_, 1000 independent runs with randomized initial conditions were conducted to ensure convergence. The covariance type was set to full, and the Expectation-Maximization (EM) algorithm was initialized using the k-means++ method. A Dirichlet process prior was applied to the component weights, enabling flexible modeling of the underlying data distribution. Convergence was assessed using a tolerance of 10^−4^, with a maximum of 5000 EM iterations allowed. All other hyper-parameters of the BGM model were kept at their default values. This clustering approach allowed for the identification of distinct populations or patterns in the data, supporting downstream analyses of translocation dynamics and signal variability across experimental conditions.

### Monitoring Amino Acid Volume Occupancy in the Pore during Molecular Dynamics

The volume of a spherical cap was used to estimate the portion of each amino acid residing inside the nanopore during its passage through the cylindrical channel. This volume corresponds to the geometric intersection between the amino acid, modeled as a sphere, and the pore, modeled as a cylinder. Each amino acid along the sequence was modeled as a sphere centered at its center of mass, with radius derived from its known total volume. The pore was modeled as a cylinder of effective diameter and thickness as given in the Methods above. The volume of the spherical cap, *V*_cap_, is given by:

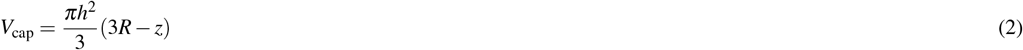

where *R* is the radius of the sphere and *z* is the height of the cap, corresponding to the portion of the amino acid inside the pore. The height is computed from the position of the amino acid center of mass along the nanopore axis (*z*-coordinate) relative to the pore top and bottom boundaries. The volume inside the pore was then computed as the difference between the spherical cap volumes as the amino acid enters and exits the pore, yielding a “slice volume” that represents the fraction of the residue physically located within the sensing region. The position of the amino acid center of mass along the nanopore axis (*z*-coordinate) was used to calculate this dynamic intersection. Finally, the volume traces obtained from MD simulations were filtered using the same parameters applied to ionic current signals (see above) to allow direct comparison across these datasets 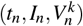, where *n* represents the frame index and *k* the amino acid index. Sequence motifs were associated to specific steps in the ionic current signal if the overlap between the volume and the step traces was larger than 60%.

## ACKNOWLEDGEMENTS

The simulations were performed using supercomputer facilities at the “Mesocentre de calcul de Bourgogne-Franche-Comté (MesoBFC)”, as part of the PARKIN-NANO project (Défi CPU 2023). In addition, the work is part of the project SEPIA supported by the EIPHI Graduate School (contract ANR-17-EURE-0002), the Conseil Régional de Bourgogne Franche-Comté and the European Union through the PO FEDER-FSE Bourgogne 2021/2027 programs.

## AUTHOR CONTRIBUTIONS

A.N. devised and planned all simulations. P.D. carried out MD simulations, and A.N., P.D. and A.U.H. performed the analyses. The results were collectively discussed. A.N. wrote the paper, incorporating inputs from all co-authors. P.S. supervised the project.

## ADDITIONAL INFORMATION

**Supplementary information** accompanies the paper: 3 figures, see PDF file and 2 movies available here: https://filesender.renater.fr/?s=download&token=2ef825c3-cb92-4099-b526-bbcd40f4f086

