## Supplementary Information for "Sequencing of *α*-synuclein Intrinsically Disordered Protein in MoS_2_ Nanopores"

### LIST OF CONTENTS

• **Figure S1:** Stepwise signal recorded during the sequencing of WT $\alpha$S, A30P and E46 mutants through single-layer $MoS_2$ nanopores.

• **Figure S2:** Statistical analysis of stepwise signal characteristics recorded during the sequencing of WT $\alpha$S, A30P and E46 mutants through single-layer $MoS_2$ nanopores.

• **Figure S3:** Volume of amino acids inside the nanopore (in Å$^3$) and corresponding ionic current traces during the sequencing of WT $\alpha$S, A30P and E46K mutants.

• **Movie S1:** Translocation of the WT $\alpha$S protein through single-layer $MoS_2$ nanopores (read 1). The timestep is 1 ns, the duration of the simulation is 1.7 $\mu$s. The color code is the following: for the membrane, Mo (density) and S (yelloworange); for the ions, $K^+$ (aquamarine) and $Cl^-$ (marine); for the peptide: Ala (lime), Arg (density), Asn (salmon), Asp (warmpink), Cys (paleyellow), Gln (tv_red), Glu (ruby), Gly (white), His (slate), Ile (forest), Leu (smudge), Lys (deepblue), Met (sand), Phe (gray40), Pro (gray20), Ser (tv_orange), Thr (brown), Trp (palegreen), Tyr (wheat), Val (pink). The images were generated using PyMOL (version 3.0.3)

• **Movie S2:** Ionic current trace recorded during the translocation of the WT $\alpha$S protein through single-layer $MoS_2$ nanopores (read 1, see Movie S1). The timing and duration is the same as in Movie S1. Amino acid inside the pore are also indicated, using the same color code as in Movie S1.

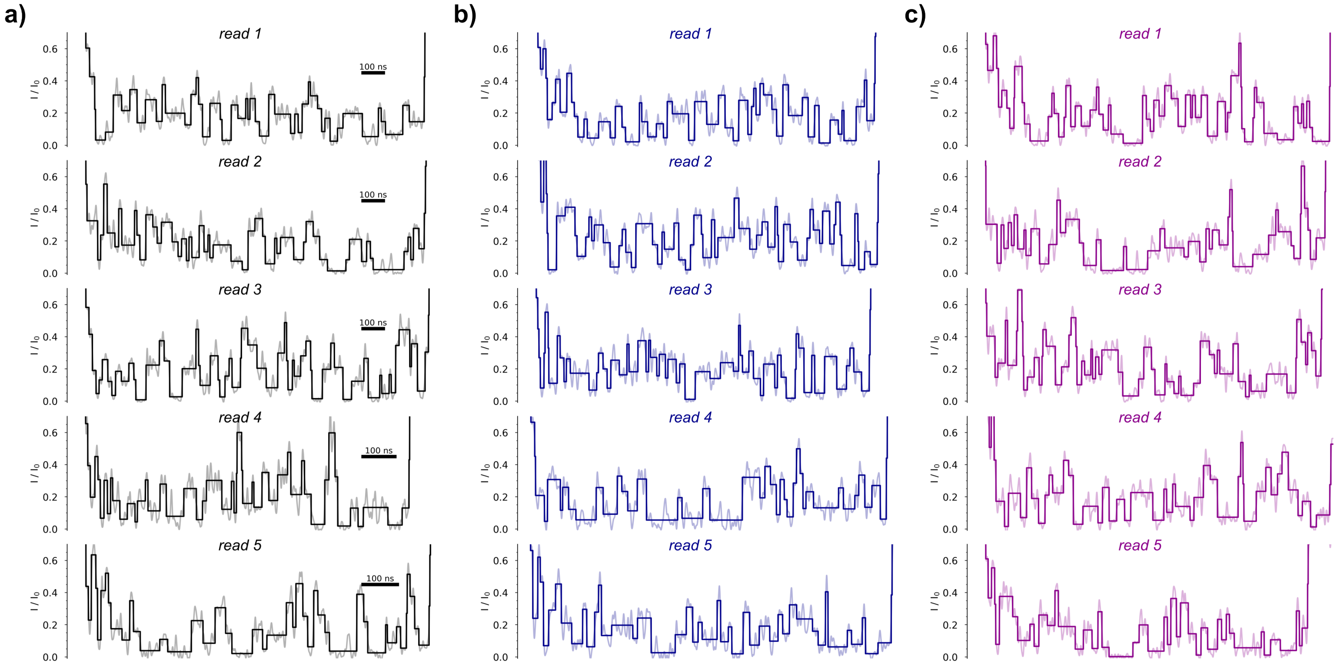


**Figure S1.** Stepwise signals derived from the ionic current traces recorded during the sequencing of WT $\alpha$S (panel a), A30P (panel b) and E46 mutant (panel c) through single-layer $MoS_2$ nanopores.

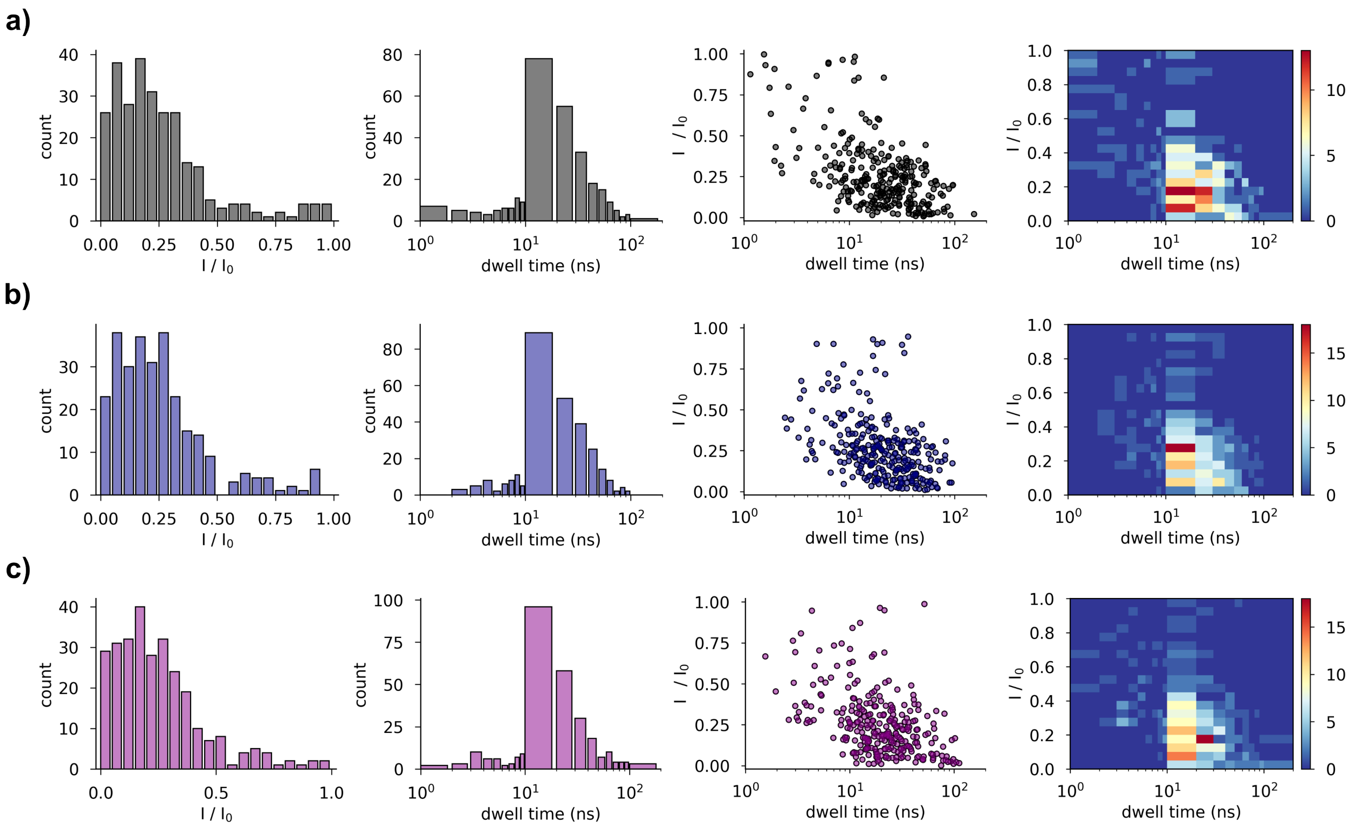


**Figure S2.** Statistical analysis of stepwise signal characteristics, *i.e.*dwell time $\tau$ (in ns) and relative depth $I/I_0$ of the current blockade, recorded during the sequencing of WT $\alpha$S (panel a), A30P (panel b) and E46 mutant (panel c) through single-layer $MoS_2$ nanopores.

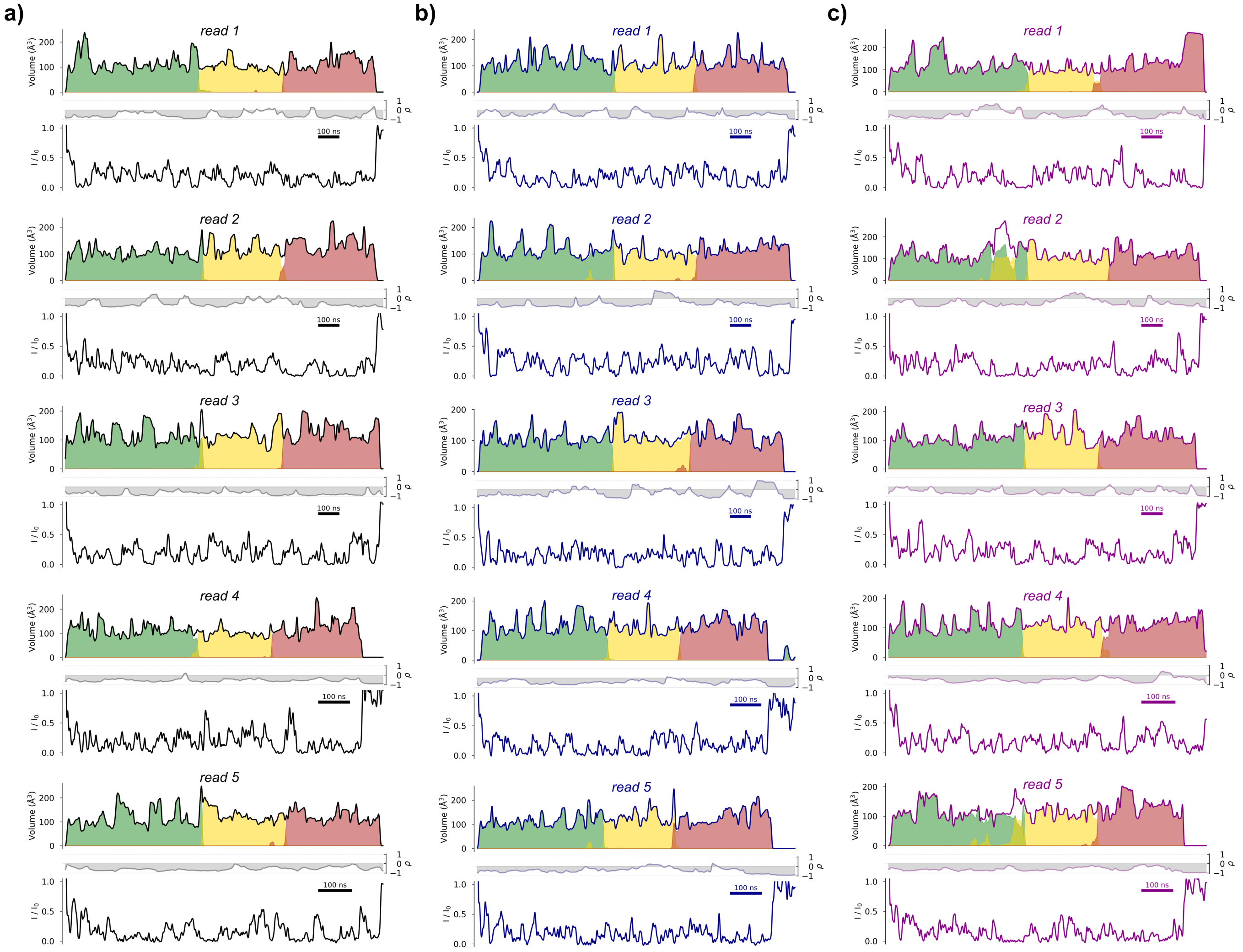


**Figure S3.** Volume of amino acids inside the nanopore (in Å$^3$) and corresponding ionic current trace during the sequencing of WT $\alpha$S (panel a), A30P (panel b) and E46K mutant (panel c). Middle panel shows the Pearson correlation $\rho$ over time.
